# Spatially-structured inflammatory response in the presence of a uniform stimulus

**DOI:** 10.1101/2025.01.28.635318

**Authors:** Elizabeth R. Jerison, Nicolas Romeo, Stephen R. Quake

## Abstract

Inflammatory responses occur within the complex spatial context of tissues and organs, and many questions remain about how tissue structure and cellular communication shape their spatiotemporal dynamics. Here, we use a multiplexed RNA in situ hybridization approach, together with analytical tools, to study inflammatory gene expression in the larval zebrafish tailfin in response to a bath of lipopolysaccharide (LPS). We use this model system to address whether spatial structure emerges in the tissue response even absent the spatial variation introduced by a pathogen. We find that epithelial cells in the tailfin express several pro-inflammatory genes, and that across these genes, the uniform stimulus triggers a spatially non-uniform response. We use a graph-based spectral decomposition method to analyze its structure, and find that long modes dominate, creating zones of activation. Overall, these zones account for a majority of the variation in gene expression. Our results show that epithelial cells are important producers of pro-inflammatory effector molecules in this system, and that tissue induces spatial correlations even absent a structured input.

---

Inflammatory responses are controlled by signaling amongst immune and other cells in tissue environments. These cells detect foreign, pathogen-derived molecules or host, damage-associated molecules, and may react by producing pro-inflammatory cytokines and small molecules (Murphy & Weaver, 2016; Medzhitov, 2010). Considerable effort has been devoted to understanding these signaling input/output relationships in cell culture systems, between stimuli such as lipopolysaccharide (LPS) and TNF, and cellular activation or gene expression (for recent reviews, see (Sheu *et al*., 2019; Son *et al*., 2023; Kizilirmak *et al*., 2022)). These studies have yielded a detailed molecular understanding of cellular decoding of inflammatory signals in many cases (Luecke *et al*., 2021; Altan-Bonnet & Mukherjee, 2019).

However, in living organisms these responses occur in the spatial context of tissues and organs. The collective response of the tissue may differ widely from that of individual cells: for example, recent work has demonstrated propagation of waves of Erk activity through tissue during regeneration (De Simone *et al*., 2021), a phenomenon that has no analog in a non-spatial setting. In the context of inflammatory and immune signaling, theoretical efforts in reaction-diffusion frameworks have demonstrated that a broad range of spatiotemporal phenomena are possible (Forsten & Lauffen-burger, 1992; Shvartsman *et al*., 2001; Coppey *et al*., 2007; Segel *et al*., 1992; Penner *et al*., 2012; Yde *et al*., 2011a; Vig & Wolgemuth, 2014). These modeling frameworks hold promise for *in silico* exploration of the collective response, and for predicting the spatial spread of inflammation under a variety of stimuli. Recently, several studies have made progress in connecting such frameworks with experiments in cultured systems of one or two cell types (Oyler-Yaniv *et al*., 2017; Bagnall *et al*., 2018; Son *et al*., 2022). These studies have demonstrated that the type and scale of the response is inextricably dependent on spatial context: for example, the scale of a cytokine niche is set by the density of consuming cells near a producing cell (Oyler-Yaniv *et al*., 2017), and whether a signaling front propagates is dependent on the density of cells participating in signal relay (Bagnall *et al*., 2018).

This work suggests that the response of the tissue may differ substantially from that of cultured individual cells. To probe these differences, it would be ideal to investigate simple but realistic model tissue systems that are amenable to controlled perturbations. Such systems would allow us to study, in a variety of contexts, how spatial cooperation and tissue structure shape inflammatory response.

In particular, since pathogens, wounds, and other immune stimuli are usually spatially non-uniform, it is not clear what, if any, role the tissue itself plays in generating a spatially-structured response. Are there spatial patterns or structure that are emergent from the tissue? Are there intrinsic length scales to the tissue response? How can we detect these given that stimuli introduce their own length scales?

Here, we use a multiplexed RNA fluorescence in situ hybridization (FISH) approach to measure spatial patterns of inflammatory signaling in the tailfin of larval zebrafish exposed to a bath of lipopolysaccharide (LPS). This system provides a model of an intact epithelial double bilayer (Eisenhoffer *et al*., 2017; Cokus *et al*., 2019) in a two-dimensional geometry, responding to a spatially-uniform stimulus. We use scRNA-seq to identify candidate response genes, and measure the spatial distribution of the expression of these genes in fixed, whole-mount tailfins. We find that cells in the epithelium express several genes for secreted pro-inflammatory factors during the response, including the apical cytokine il1b, metalloproteases, and prostaglandins. Across these genes, the spatially uniform stimulus elicits a spatially non-uniform response. We develop a graph-based spectral decomposition tool to analyze contributions to the response from different length scales. This allows us to detect activated domains, and to quantify the extent to which these tissue-level effects determine the gene expression response.

## Results

### Measuring spatial patterns of gene activation during a model inflammatory response

To induce inflammation in tissue, we used an established bath immersion model of larval zebrafish in lipopolysaccharide (LPS) (Watzke *et al*., 2007; Yang *et al*., 2014; Philip *et al*., 2017). This treatment has been shown to cause a systemic inflammatory response similar to the acute phase of sepsis, including vascular leakage and tissue damage, especially in the tailfin (Philip *et al*., 2017). To identify candidate genes involved in the tailfin inflammatory response, we performed single-cell RNA sequencing on dissociated 6 dpf zebrafish larvae exposed to a bath of LPS in E3 embryo media for 10 hours, compared to vehicle control larvae (exposed to E3 embryo media alone). We focused on epithelial cells. The top 10 genes upregulated in the LPS condition (Methods) included several that are canonically associated with inflammatory response, including il1b, homologous to the apical cytokine IL-1,*B*; ptgs2b, homologous to COX2, the rate-limiting enzyme for prostaglandin synthesis that is the target of NSAIDs (Flower, 2003); and the matrix metalloproteases mmp9 and mmp13a (Manicone & McGuire, 2008). We chose 7 of the top 9 genes as candidates for spatial expression analysis, excluding 1 long-noncoding RNA and the tropomyosin tpm4a. We note that 5 of these genes are either secreted inflammatory factors or involved in synthesizing secreted factors: il1b; ptgs2b (prostaglandin synthesis); acsl4b (eicosanoid synthesis (Kuwata & Hara, 2019)); and mmp9 and mmp13a. We added to this panel the macrophage marker mpeg1.1 (Ellett *et al*., 2011), and the inflammatory repressor socs3a (Carow & Rottenberg, 2014; Veneman *et al*., 2013).

To measure the spatial expression patterns of these candidate response genes, we exposed 5 dpf zebrafish larvae to baths of LPS at 4 concentrations, in addition to vehicle control, for 10 hours. We also included a 4 hour timepoint for the highest LPS concentration. We note that the treatment can cause substantial morbidity via systemic inflammation (see Methods for endpoint criteria). At the specified timepoints, we fixed groups of larvae. We mounted whole tailfins on slides and used an RNAscope^*TM*^ -based smRNA FISH assay to label 9 genes in each sample, for 4-6 biological replicates per group (Methods). We measured the spatial distribution of the fluorescence intensity associated with each gene probe using laser-scanning confocal microscopy, across 3 rounds of detection for each sample. We used a DAPI stain of nuclei to cross-register the samples across imaging rounds, and performed an approximate cell segmentation using the reference DAPI (nuclear) image. We measured gene expression in two ways: via detecting spots associated with each labeled RNA, and by measuring fluorescence intensity within each segmented cell region (Methods; note that the autofluorescent tailstem region was masked). We used intensity as the primary measure for gene expression because it is valid in high expression regions, where spots lie in too close spatial proximity to be counted; however, the conclusions presented do not depend on which measure is used (see SI for parallel figures using spot counts).

We observed expression of the candidate genes in the LPS-treated tailfins (Figure 1, Figure S1, Figure S2). The macrophage marker, mpeg1.1, was present in isolated clusters consistent with sparse macrophages, and showed no trend across treatment conditions (Figure S2). For the remaining 8 genes, the response was dose-dependent, only weakly deviating from control at 20 *μg/mL*, and strong at 30 *μg/mL*, driven by a tail of high-expressing cells at the lower concentrations (Figure 1B, Figure S1). We note that in 1/6 vehicle control samples, we observed some clear expression of mmp13a, mmp9, odc1, and il1b, which may be due to a damage response in this tailfin. Interestingly, socs3a, which was chosen on the basis of the literature rather than via the scRNAseq screen, showed increased expression in the highly activated samples (Figure 1B). These observations show that during the inflammatory response accompanying LPS exposure, tailfin epithelial cells express genes related to production of a number of pro-inflammatory factors, including il1b, prostaglandins, and metalloproteases.

**Figure 1:**
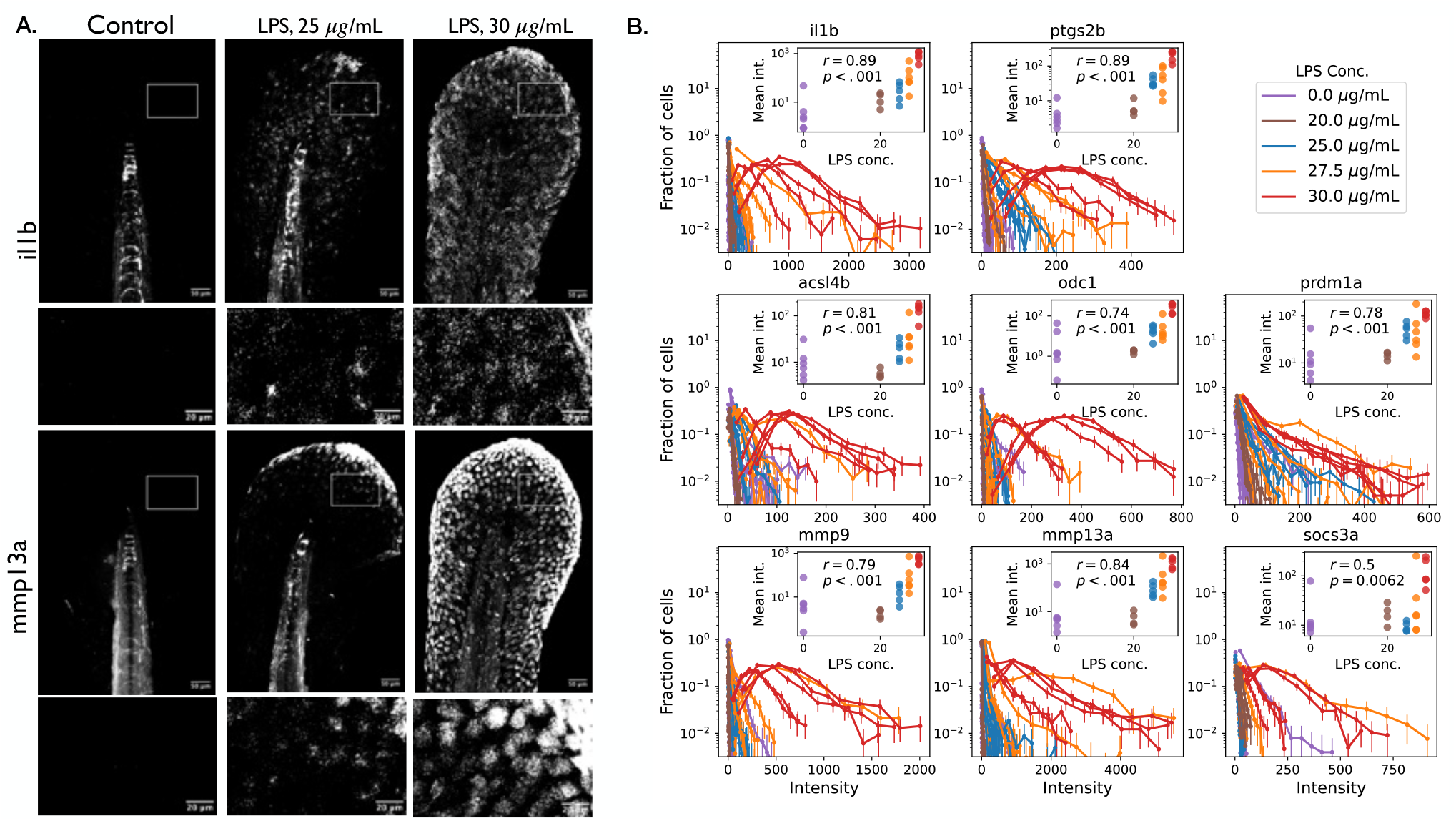
Inflammatory gene expression response to LPS. A. Examples of *in situ* images for Control, 25 *μg/mL*, and 30 *μg/mL* treatments, for the il1b and mmp13a fluorescence channels. Each column is a tailfin sample. Maximum Z projection of tiled Z stacks. B. Distribution of intensity per cell region for candidate response gene channels across all samples. Distributions represent histograms with equal-width bins up to the 98th percentile. Error bars: 95% confidence intervals on a bootstrap over cell regions. Inset: mean intensities per sample as a function of LPS concentration (log scale). Spearman rank correlation coefficient and p value under a permutation test.

### A heterogeneous, spatially structured response

Since we observed minimal activation at 20 *μg/mL*, for this and subsequent sections, we analyzed the concentration groups with a medium or high response–25, 27.5, and 30 *μg/mL*. When presented with a uniform stimulus, all cells could turn on a common set of genes at similar levels. This simplest null model would preclude any spatial structure in the response. In contrast, variability is a visually prominent feature of the in situ images (Figure 1A). Indeed, across nearly all genes and samples, the true expression distribution is far broader than a null based on re-distributing spots, accounting for potential variation in true number of cells per segmented cell region (Figure 2A, Figure S3). In particular, there is a heavy tail of high-expressing cells that are inconsistent with the uniform expression model (see Figure 2A, inset for an example). This variability is consistent with results from cell culture experiments, which have showed that individual cellular responses, even to a constant, uniform stimulus, are often highly heterogeneous (Tay *et al*., 2010; Singer *et al*., 2014; Kramer *et al*., 2022).

**Figure 2:**
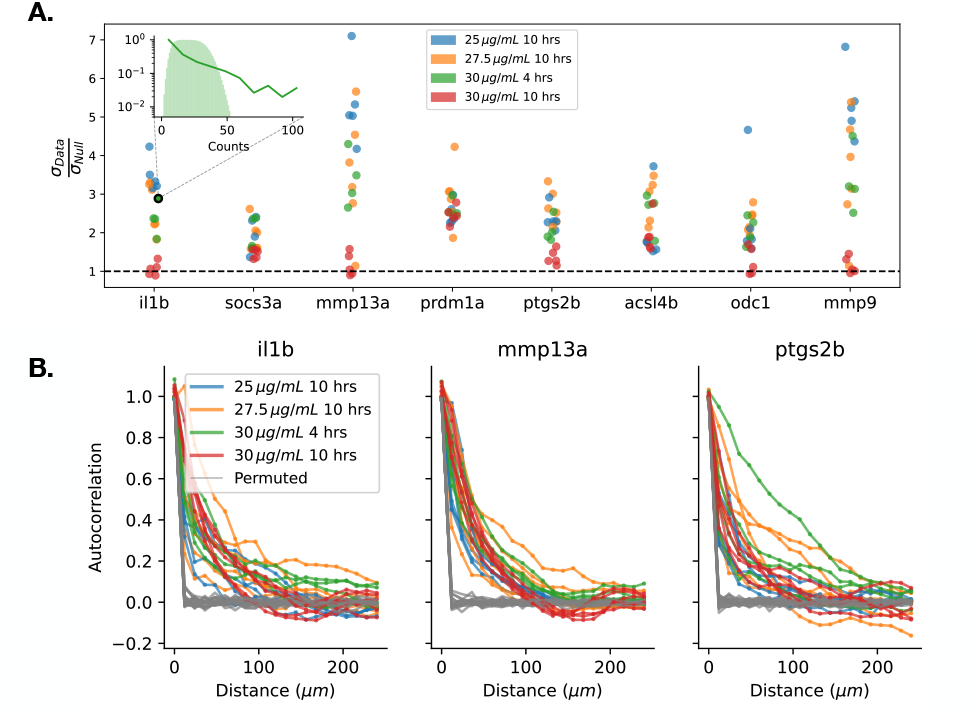
Cells activate non-uniformly. A. Width of counts distribution compared to a null distribution modeling random redistribution of spots. The null distribution was constructed assuming a uniform distribution of .5 to 2.5 true cells per segmented cell region (Methods); see Figure S3 for additional choices. Inset: example of counts distribution and null. B. Spatial autocorrelation of gene expression for 3 example genes. Grey curves represent a permutation on cell region locations in each sample. See Figures S4, S5 for all genes.

Our main goal is to understand whether and how spatial structure contributes to this heterogeneity. To begin addressing this question, we assessed whether highly-activated cells were randomly distributed. To do so, we measured the spatial autocorrelation of expression of each gene. We found that there is substantial autocorrelation for all genes compared with permuted samples, with deviations ranging out towards the full system size (Figure 2B, Figure S4).

### Anatomy biases but does not fully explain activation

As observed above, gene expression across cells is heterogeneous, and this heterogeneity is not randomly organized in space. This shows that there is some role of tissue structure in determining the response. There are two main, not mutually exclusive, ways in which spatial correlations could arise in tissue. First, spatially-varying gene activation could be driven by anatomy. In this case, as in developmental contexts, gene expression roles would vary based on location, leading to pre-patterned zones of activation for different genes. Alternatively, zones of activation could arise spontaneously at any location due to secondary signaling and biophysical cues.

To investigate the role of anatomy in determining gene expression patterns, we sought to map tailfin samples to a standard set of coordinates. In our system, the most relevant anatomical coordinates are the distance from the tail edge (outer vs. inner), and the axial location. To assess evidence for patterning with respect to these coordinates, we used a 2D mesh (Coons (Coons, 1967)) parametrization to unfold each tail sample, mapping it to a rectangle (Figure 3A). For this analysis, we excluded 4 samples where part of the tail edge folded over during mounting (see Figure S6 for all sample outlines with the excluded samples indicated).

**Figure 3:**
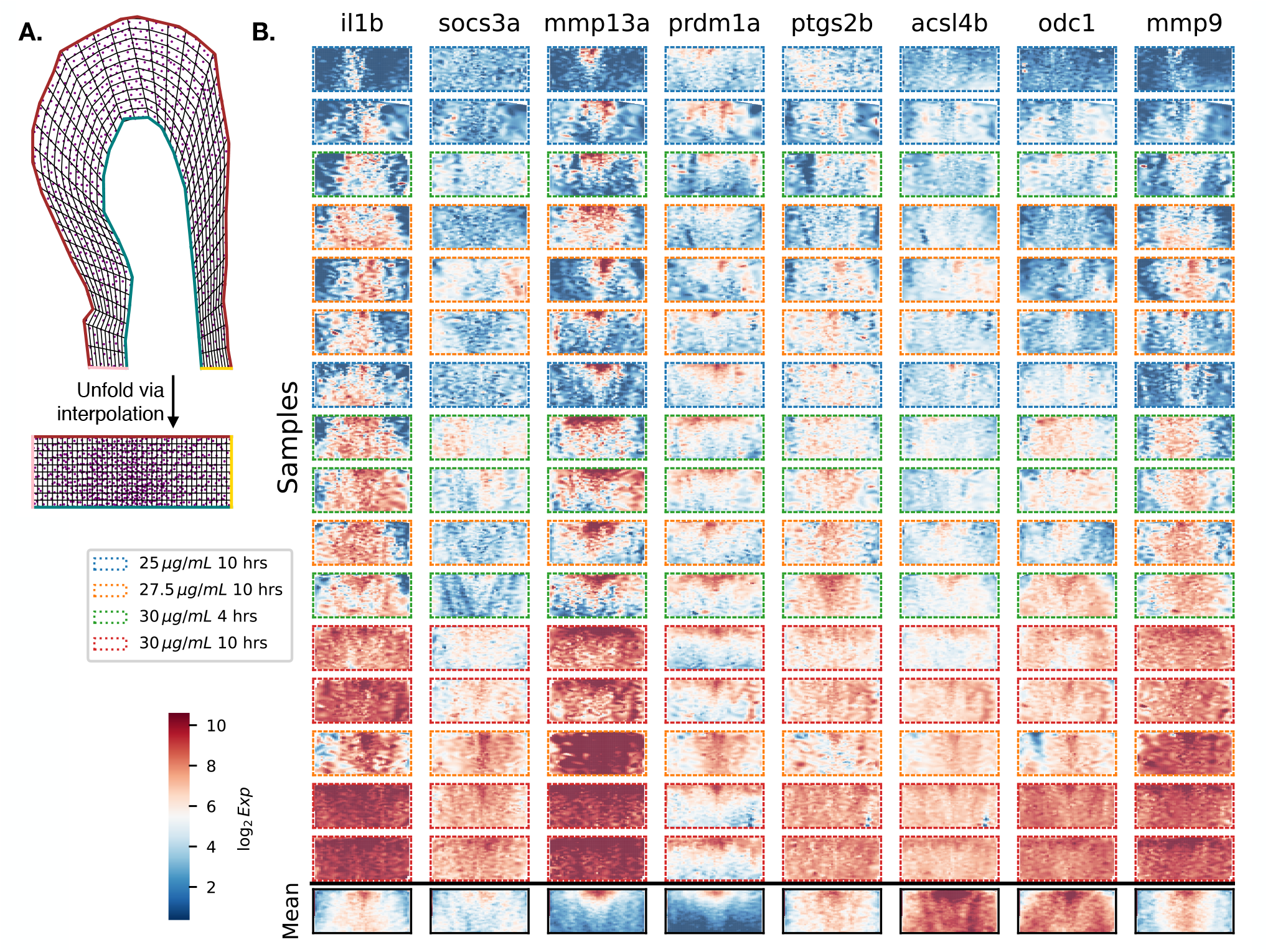
Anatomical biases and variable activated regions across samples and genes. A. Example of bilinear (Coons) interpolation to map a tail region to a rectangle. Dots show locations of cell region centroids. B. Heat maps of expression of the response genes (log scale), in all samples passing the morphology filter, ordered from lowest to highest total expression. Expression data has been interpolated onto grid points within the rectangle mapping; colormap ranges from the 2nd to 98th percentile of expression values across all genes and samples. Outline color indicates the treatment group. Final row: mean of expression across samples for each gene (log scale, normalized by standard deviation within each sample. Colormap ranges from the 2nd to the 98th percentile across these means).

Ordering samples from low to high overall activation amplitude (Figure 3B, Figure S7), we observe that at the highest activation levels, cells throughout the sample express high levels of all gene markers except prdm1a, which remains generally localized to the top (outer) edge. Thus cells throughout the sample can in principle express all of these genes; there are not anatomical zones that exclusively turn on different genes. Further, high activation regions for each gene vary between tailfin samples (Figure 3B). This supports spontaneous chemical and/or biophysical inflamed domains, which may spread during the response, instead of tissue pre-patterning.

However, there are clear anatomical biases to the activation pattern, as shown by the average expression pattern across all samples (Figure 3B, bottom row). In particular, mmp13a, prdm1a, and mmp9 have distinct spatial organization.

Combined, these observations indicate that anatomy biases but does not fully explain activation: cells in certain anatomical regions are more likely to activate, but do not do so deterministically.

Surprisingly, the large-scale spatial activation patterns, both on average (Figure 3, bottom row) and within each sample (Figure 3, rows), are distinct from gene to gene. This indicates that high response zones cannot be accounted for only by an inhomogeneous distribution of LPS within tissue, which would create only 1 large-scale spatial pattern per tailfin. Rather, the observed response patterns are due to a combination of pre-patterned tissue biases, fields of secondary signaling molecules, and biophysical cues.

### Length-scales of the response via spectral graph decomposition

Since anatomy does not fully explain the spatial organization of the response, we sought a more flexible tool to dissect its structure. In particular, we would like a way to detect and quantify domains of activation, even if they do not arise in stereotyped locations within each sample. More generally, we would like to ask about the structure of the response: Is it scale-free, or are there dominant length-scales? Does structure arise primarily at small scales, such that it averages out over larger domains, generating a collectively uniform response at the macroscopic level? Or, alternatively, are there larger-scale spatial domains that remain after averaging at smaller scales, even in cases without clear anatomical patterning?

To address these questions, we would like a method for quantifying and separating contributions from different spatial scales (Romeo *et al*., 2021). The standard approach to such a problem uses Fourier decomposition to measure the power attributable to each spatial mode. Such decompositions are useful for measuring contributions from long- and short-wavelength behavior; for detecting spatial structure at particular intermediate length scales; and for filtering short wavelength noise. However, most such methods are not suited to small, irregularly shaped regions, which preclude their use with our and many other biological samples.

Here we use signal graph processing (Ortega *et al*., 2018), a graph-based spectral approach: we represent each sample as a graph, where each cell is a node and adjacent cells are connected by edges. We can then compute the eigenvectors of the normalized graph Laplacian; this set of orthonormal basis vectors allows the definition of an analog of the Fourier decomposition that captures large- to small-scale spatial structure in a manner that respects the geometry of the sample (Figure 4A). Vectors of gene expression values can be decomposed onto these spatial modes to isolate long- and short-wavelength variation. We note that there are the same number of modes as cells in the sample; including all modes allows for an exact reconstruction of the signal, and a projection onto a subset of modes represents an approximate reconstruction.

**Figure 4:**
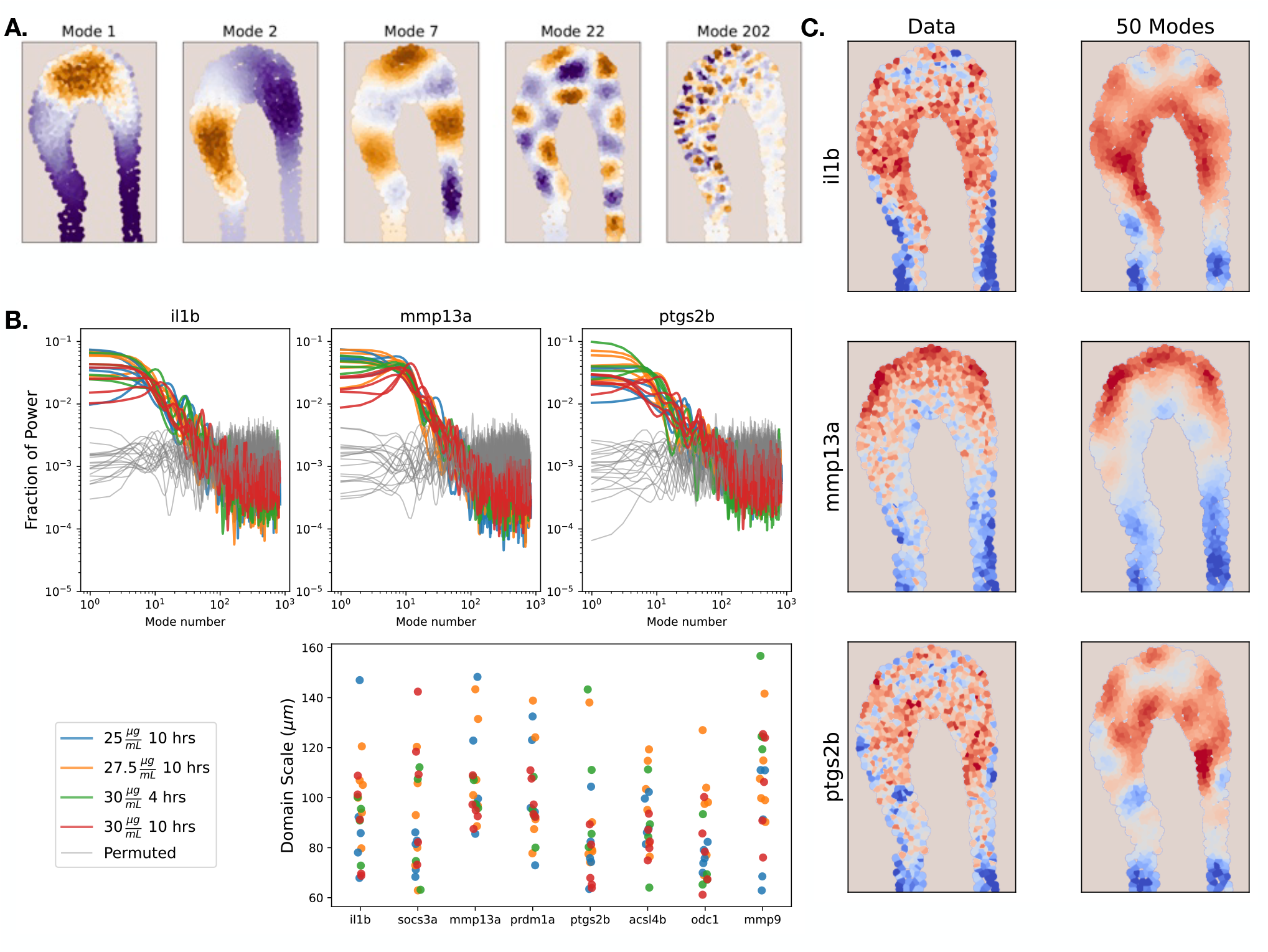
Extraction of length scales and long modes via spectral graph decomposition. A. Heatmap visualization of graph eigenvectors for one sample, where the color represents the magnitude and sign for the eigenvector component for each cell. Low modes capture long-wavelength variation, and high modes capture short wavelength variation. B. Top row: Power as a function of mode number, for 3 example genes (see Figures S8, S9 for all genes). Spectra have been smoothed with a Gaussian of width *CJ* = 3 modes. The spectra show domains at large spatial scales, transitioning to unstructured fluctuations. Bottom: Correlation length scale (domain scale) for each sample and gene, determined based on the mean absolute deviation of the power spectra (Methods). C. Examples of gene-by-gene reconstruction of the data using the first 50 graph eigenmodes, representing a spatial long-pass filter.

Applying this decomposition to our samples, we observe that the spectra do not show scale free (power law) behavior (Figures 4B, S8, S9); rather, there is a transition from larger-scale correlated domains to uncorrelated fluctuations. We used the mean absolute deviation of the power spectrum to measure the scale of the plateau (Methods), and extracted from this spatial frequency a characteristic domain size for each sample and gene (Figure 4B, second row). These are typically about 100 *μm*, or 10 cell lengths.

From the decomposition, we can extract large scale spatial patterns without regard to location within the sample, via projection onto a subset of long modes (Figure 4C, Figure S10). The reconstruction acts as a long-pass filter, preserving the large-scale structure of gene expression while suppressing cell-to-cell fluctuations.

### The majority of variation in gene expression is explained by spatial effects

The spectral decomposition provides a natural way to quantify the overall contribution of spatial structure to explaining the variability in gene expression response. In particular, the cumulative fraction of the power in a set of modes corresponds to the fraction of the variance that is due to these modes. Here, we are interested in the amount of variability explained by long modes, which gives us a measure of the importance of tissue-level effects, as opposed to cell-to-cell fluctuations.

From the cumulative fraction of the variance as a function of mode number (Figure 5A, Figure S11), we observe that there is a turnover (corresponding to the plateau location in the spectra, Figure 4B, Figure S11). We determine the location of this turnover by measuring the mode at which the power first falls below the noise floor, and calculate the fraction of variance attributable to the longer modes.

**Figure 5:**
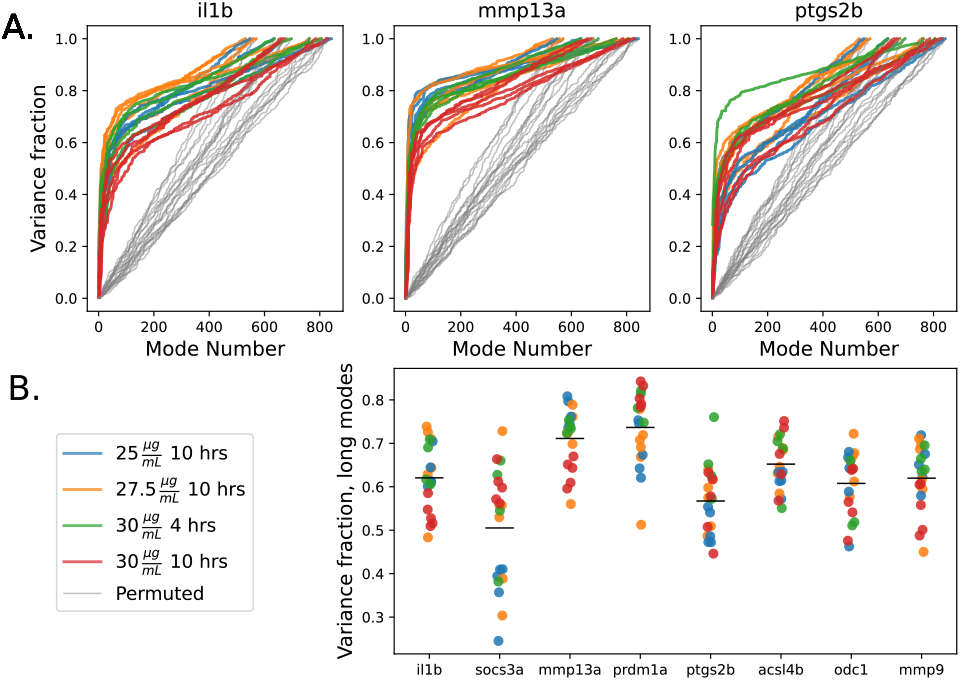
The majority of variance is explained by long spatial modes. A. Fraction of the variance explained by modes up to the given number (see Figure S11 for all genes). B. Fraction of the variance explained by the longer modes, defined based on the location of the slope change in the cumulative power, as seen in A. Black bars represent the average over all samples for each gene.

In each case (Figure 5B, Figure S12), the long spatial modes capture a large fraction of the variance. Note that the effect is the weakest for socs3a, which, as discussed above, shows strong activation only in the highest response samples. For the other genes, these long modes typically account for a majority of the variance in gene expression. This shows that tissue-level effects are integral to determining the cellular response.

As noted, in our system, these effects arise both due to inherent anatomical bias and to non-stereotyped domains of about 100 *μm* in scale. Combined, these observations show that the tissue contributes to the structure of the response both through pre-patterning and spontaneous correlated zones.

## Discussion

The advent of single-molecule in situ hybridization techniques, which are increasingly amenable to multiplexing, as well as in situ RNA sequencing techniques, has enabled enormous growth in measurements of spatial patterns of gene expression in tissue. These techniques provide a new set of tools for uncovering influences of space and tissue context on inflammatory response. Recent studies have begun deploying these techniques extensively in complex clinical settings (Gniadecki *et al*., 2023; Schäbitz *et al*., 2022; Caetano *et al*., 2023; Son *et al*., 2022).

By contrast, there are very few studies that measure response of inflammatory genes in tissue to a known stimulus. Such studies are necessary if we hope to disentangle the role of the tissue vs. the stimulus in driving inflammatory response.

Here, we used a bath of LPS to stimulate inflammation in the larval zebrafish, and studied gene expression response in the tailfin. We found that this spatially uniform stimulus elicits a spatially non-uniform response, characterized by 100 *μm* scale spatial domains, as well as additional uncorrelated, fine-scale heterogeneity.

Cytokine signaling is usually considered to be the purview of professional immune cells, which are sparse in most tissues. The extent to which these cells, vs all of the cells in the continuum, participate in detection of pathogen and damage cues and secretion of cytokines, is an important factor in understanding the spatial organization and control of such a response. Our data provides evidence that zebrafish tailfin epithelial cells express il1b, an important pro-inflammatory cytokine, during the LPS response. This is consistent with prior work showing that zebrafish tailfin epithelial cells express il1b in response to injury (Hasegawa *et al*., 2017). Additionally, activation of NF-*κ*B and Il1b expression has been observed in mouse airway epithelial cells in response to inhaled LPS (Skerrett *et al*., 2004). More broadly, epithelial cells in mucosal barrier tissues such as the lung and gut are important sources of cytokines (Whitsett & Alenghat, 2015; Peterson & Artis, 2014). Our data thus contribute to the body of evidence that many cells in the continuum of barrier tissues, not only sparse professional immune cells, participate in the establishment and propagation of cytokine signaling. We also observe gene expression related to prostaglandin synthesis and matrix metalloprotease production, processes commonly associated with inflammatory response in epithelial cells (Whitsett & Alenghat, 2015).

The presence of large-scale spatial domains in our data could be due to two (not mutually exclusive) effects. First, it is possible that fine variation in cell state or type correlate with spatial location in the tail and lead to physiologically-biased response zones. Second, it is possible that the cells are responding to fields of secondary signalling molecules and biophysical cues generated within the system, whose distribution is spatially non-uniform. Our data support a role for both of these effects: there are anatomical biases on average, but also zones of activation at non-stereotyped locations. In both cases, the presence of these large-scale domains indicates that tissue structure is a primary determinant of the response.

Since we labeled multiple response genes simultaneously, we were able to observe that the spatial response pattern differed from gene to gene within each sample. This indicates that high response zones cannot be accounted for only by an inhomogeneous distribution of LPS within tissue, which would create only 1 large-scale spatial pattern per tailfin. Rather, it again suggests that multiple different secondary signaling molecules and/or biophysical cues combine to elicit expression of different genes.

To separate long and short-wavelength variation, we have used a graph-based spectral decomposition approach. This computationally simple method allows application of standard wavelength-decomposition tools to samples of any shape, with masked regions and irregular boundaries, and thus may be a useful addition to the suite of approaches recently developed for detecting spatial patterns in multiplexed FISH and spatial transcriptomic data (Tian *et al*., 2023; Walker *et al*., 2022).

Our data indicate that, in this system, cells respond heterogeneously to the inflammatory environment, and that this heterogeneity cannot simply be locally averaged away, but rather generates larger-scale biases in gene expression. The spectral decomposition allows us to quantify the contribution of these tissue-level effects to the response. We find that they account for a majority of the variance in gene expression in this system. Additional cell-to-cell variability is also present, and is a slightly less important contributor. Thus there is spontaneous generation of activated zones for each gene within the tissue.

What are the implications of this spatial heterogeneity for control of inflammation in tissue? The consequences of spatial heterogeneity for immune signaling have been studied most intensively for ‘sender-receiver’ systems, where immune cells secrete cytokines that are consumed by surrounding cells (Altan-Bonnet & Mukherjee, 2019; Oyler-Yaniv *et al*., 2017; Centofanti *et al*., 2023). In this setting, static cytokine niches with length scales from tens to hundreds of microns form around producing cells. In our system, similar patterns could arise for the secreted factors studied here, including the cytokine il1b and the metalloproteases mmp9 and mmp13a, generating local gradients around activated zones due to passive consumption by nearby cells.

However, the epithelial inflammatory process studied here differs from sender-receiver systems in two important ways, which could drive more dramatic spatial effects. First, cells throughout the tissue layer are both senders and receivers; and, second, the pro-inflammatory factors can generate positive feedback. For example, secreted il1b can activate NF-*κ*B in neighboring cells, driving transcription of itself and numerous additional pro-inflammatory signaling factors. Reaction-diffusion modeling has shown that this positive feedback can generate spreading waves and fronts in addition to stable niches (Yde *et al*., 2011b). Thus, activated zones of these factors could drive runaway responses in tissue. An inherent limitation of the RNA in situ approach used here is that it captures snapshots in time, so that it is not possible to directly probe such dynamical processes. Live microscopic investigation of the formation and spread of inflammatory domains in tissue would be an interesting avenue for future work.

## Materials and Methods

### Zebrafish and husbandry

Adult AB (wildtype) zebrafish were purchased from the Zebrafish International Resource Center (Zirc) and housed at 28 C. Zebrafish were mated using standard husbandry practices to generate embryos. All procedures were approved by the IACUC Animal Care and Use Committee of Stanford University.

### scRNAseq screen for candidate genes

6 dpf AB zebrafish larvae were immersed in either E3 with 37.5 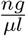 LPS, or E3 alone (vehicle control), for 9.5 hours at room temperature. 12 larvae per treatment group were then transferred to .04% w/v Tricaine, and decapitated with a sterile razor blade. Tail regions from the individuals in each treatment group were pooled and dissociated; see Supplementary Information for a detailed dissociation protocol. The 10X Genomics Chromium 3’ platform and reagents (v3.1 kit) was used for cell encapsulation, reverse transcription, and library preparation, according to manufacturer instructions (manual revision D), with 12 cycles of preamplification. For cell encapsulation, one lane of the 10X chip was used for each treatment group (LPS and control), at a target of 16,000 cells/lane. Libraries were sequenced on a NovaSeq 6000, following 10X manual instructions, with a target of 50000 reads/cell. Sequencing data was aligned to the zebrafish reference genome (GRCz11) using 10X cellranger software (6.1.2).

To identify candidate genes that activated in epithelial tissue, the software packages decontX, scrublet (run in RStudio), and scanpy (run in Python v3.7) were used to generate quality scores for cells; cells with > 12% mitochondrial reads, scrublet score > .3, or decontX score > .5, were eliminated, as well as genes expressed in *<* 10 cells. The Leiden algorithm in scanpy was used to cluster cells, and a basal epithelial cell cluster in each sample was identified based on high expression of the marker genes tp63, mmp30, cxcl8a, and egfl6. To filter genes that might arise due to differences in ambient RNA background in the two 10X lanes, a cluster of red blood cells was identified within each sample, and a list of genes differentially expressed between the LPS and Control red blood cell clusters was determined via a Mann-Whitney U test; these were considered false detections. To generate a candidate gene panel for in situ labeling, the 90th percentile of expression of each gene in LPS vs. control were compared, any genes in the red blood cell false detection list were filtered, and the remaining top genes were identified.

### Zebrafish LPS exposure and fixation

Zebrafish larvae at 5 dpf were immersed in 2 mL E3 (Westerfield, 2007) (vehicle control), or 2 mL E3 supplemented with 20, 25, 27.5, or 30 *μg/mL* Lipopolysaccharide (LPS) from *Pseuodomonas aeruginosa* (Sigma L9143, batch 0000085561). The experimental endpoint was reached if larval fish were unresponsive to gentle squirting with a pipette. Exposures were carried out at room temperature for 10 hours for all conditions, and additionally for 4 hours for the 30 *μg/mL* concentration group. The proportion of fish in each treatment group that was removed prior to the collection timepoint in each treatment group was: Vehicle Control (0), 20 *μg/mL* (0), 25 *μg/mL* (13%), 27.5 *μg/mL* (13%), 30 *μg/mL*, 4 hours (0), 30 *μg/mL*, 10 hours (35%).

At the collection timepoint, larval fish were euthanized via a 5 minute exposure to an overdose of pre-chilled Tricaine, and transferred to 4% w/v PFA in PBS to fix. Fixation was performed in 2 ml eppendorf tubes, with 5 larvae/tube, at 4 C, for 18 hours, under gentle rotation. After fixation, samples were transferred to a 33% w/v sucrose solution for 4 hours. All but about 50 ul sucrose was removed from the tubes, and the tubes were flash-frozen on crushed dry ice, and transferred to -80 C for storage.

### RNAscope^*TM*^ multiplexed RNA fluorescence in situ hybridization

Larval fish were removed from sucrose and rinsed in sterile PBS. Tail fins were cut off using a sterile scalpel and mounted on Fisher Frost + slides. As pretreatment prior to probe application, slides were dried at room temperature for 1-2 hours, baked in an incubator at 60 C for 1 hour; post-fixed in 10% NBF (Neutral Buffered Formalin) at 4 C for 15 minutes; and dehydrated with an ethanol series (50%, 75%, 100%, 100%, 5 minutes per stage). Samples were digested with RNAscope Protease III for 30 minutes at 40 C. Subsequent steps of the RNAscope HiPlex protocol were carried out according to manufacturer instructions. Probe catalogue numbers are listed supplementary file 1. To determine background, two additional samples were labeled with a negative control probe against the gene DapB, which is not present in the zebrafish transcriptome, in every channel. Prior to imaging, samples were mounted in ProLong Gold Antifade (Thermo P36934) and covered with a # 1.5 coverslip.

Imaging was performed on a Zeiss LSM980 laser scanning confocal microscope, using AiryScan 2, for DAPI, FITC, Cy3, and Cy5 channels. Imaging was performed over tiled regions, at 25X magnification/.8 NA (oil immersion), with 1 *μm* slices. Laser intensity, gain, and other microscope settings were fixed for each channel across samples, such that intensity did not saturate in probe channels (see supplementary file 1 for settings). The 2D AiryScan processing routine in Zeiss Zen Blue software was used to generate final images.

Imaging was carried out over 3 rounds. After each imaging round, the coverslip was removed and probes were cleaved according to RNAscope Hiplex manufacturer instructions. All samples that remained intact on the slide through pretreatment and all three rounds of imaging were included for analysis.

### Image registration, segmentation, and expression measurements

Image processing was performed using custom Python code, primarily with skimage libraries (code is available at www.github.com/erjerison/spatialLPS). Images were stitched using recorded stage locations (no blending). To cross-register channels across imaging rounds, the DAPI (nuclear stain) images from rounds 2 and 3 of imaging were each cross-registered to the DAPI images from round 1. A small rigid rotation and translation were allowed in the cross-registration. A maximum Z projection of the DAPI image stacks was taken, and the rotation angle was extracted via a cross-correlation of the polar transformation of the DAPI images. A translational shift was then extracted via a cross-correlation of the rotated images. Gene probe channels from imaging rounds 2 and 3 were then rotated and translated to align with round 1. Intensities from each gene probe channel were projected into 2 dimensions via a Maximum Z projection.

For each sample, a mask for the outline of the sample was drawn manually in ImageJ (Fiji), using the maximum Z projection of the DAPI image from imaging round 1. These masks exclude the autofluorescent tail stem region. Within this outline, nuclei were located by convolving a disk with the DAPI image, and using the peak local max function in skimage.feature to find peaks within this image. Peaks were then expanded into labeled cell regions via the expand labels function of skimage.segmentation, which expands labeled regions simultaneously without generating overlap, with a distance cutoff of 100 pixels (12 *μm*) in each direction.

To measure expression level of each gene within each cell region, we thresholded the maximum Z image from each channel below using a background threshold determined based on the DapB negative control samples (the average of the 99.5th percentile of pixel intensity for each channel). We measured spot number via the peak local max function in skimage.feature, with peaks required to have pixel intensity at least 200 above the background threshold. Since nearby spots may be underdetected, we also measured the total intensity above background of each gene channel in each cell region.

For the example in situ images (Figure 1A), maximum Z images for the il1b and mmp13a channels were stitched in ImageJ (no blending). Images were thresholded below and above using fixed pixel thresholds for each channel across all samples.

### Dose response and variability analysis

Computations were performed using custom Python code (source data and code to generate each figure is available at www.github.com/erjerison/spatialLPS). Distributions of counts per cell or intensities per cell were computed as histograms of the fraction of cells at each expression level, on equal-width bins up to the 98th percentile of expression, for each sample from the 10 hour timepoint. Error bars represent the 2.5th and 97.5th percentile of the histogram value on a bootstrap over cells in each sample. Dose-response was measured as the Spearman correlation coefficient between the LPS concentration and the average intensities in each sample; p-values were computed via a 1-sided permutation test.

To assess variability in expression levels between cells, we constructed a null model of uniform expression per cell in each sample and gene channel, in the presence of finite numbers of spots and uncertainty in the number of true cells per cell region (see Supplementary Information). We computed the ratio of the standard deviation of expression level in the data and null as a measure of the degree of additional variability in the data. To compute the spatial autocorrelation for each sample and gene, we centered the data by subtracting the mean and normalized by the standard deviation. We then computed the product of the expression values between all pairs of cells, binned pairs based on inter-cell distance, and calculated the average of the product within each bin. For each gene and sample, we performed the same computation with cell labels permuted.

### Tail unfolding via bilinear (Coons) interpolation

Coordinates within each tail sample outline were mapped to a rectangle via a bilinearly blended Coons patch, which interpolates any patch with 4 boundaries to a unit square (see Supplementary Information). Since this analysis requires 4 nearly convex boundaries, and also seeks to compare morphological features, we omitted 3 samples with a folded region at the edge and 1 sample lacking 4 well-defined boundary curves (Figure S6). A cubic interpolation was used on the log of expression for each gene and sample to map to a grid. To extract average patterns across samples, grid expression values for each gene and sample were normalized by their standard deviation and averaged for each gene.

### Graph signal processing analysis

To understand spatial variation between neighboring cells across the tailfin, we perform an analog of the Fourier decomposition of the signal by representing each sample as an unweighted graph where each cell is a node and adjacent cells are connected by edges. To find the graph eigenvectors for each tail sample, we performed the eigendecomposition of the associated normalized graph Laplacian (see Supplementary Information). Letting the graph Laplacian eigenvectors be 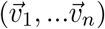, and 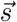 be a gene expression vector across cells in the sample, the power in each mode was computed as 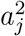, where 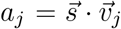. We performed the analysis on the log of the expression values, with means subtracted prior to projection. For each sample, the computation was repeated with cell labels permuted. Power spectra are shown smoothed with a Gaussian kernel with width *σ* = 3 modes. The projection onto the first 50 modes was computed as 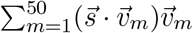.

To extract the domain scale, the noise floor of the power spectrum was computed as the average power per mode in the permuted sample. The power spectrum was then smoothed with a Gaussian kernel with width *σ* = 10 modes, and the first mode *k*^∗^ with power below the noise floor was determined. The plateau scale 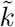 was then computed as 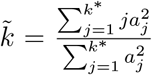. (Note that only modes up to the noise floor were included to avoid spurious contributions from the several hundred modes beyond the noise floor.) This mode scale was converted to a length scale by estimating the cellular area associated with this mode number via 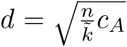, where *n* is the total number of segmented cell regions in the sample and *c*_*A*_ is the average area of one cell region, in *μm*^2^.

The variance fraction for modes up to *m* was computed as 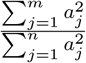, where *n* is the total number of cell regions in the sample. The variance fraction in long modes was computed as the fraction in modes up through the mode *k*^∗^, as defined above.

## Acknowledgements

We gratefully acknowledge discussions with Carlos Floyd, Ming Han, Aaron Dinner, Peter Lu, and Arvind Murugan. We acknowledge animal husbandry support from the Stanford VSC, computational resources from Stanford RCC, and the Stanford Neuroscience imaging core facility. This work was supported by the Burroughs Welcomme Foundation through a Career Award at the Scientific Interface (ERJ), the Chan Zuckerberg Initiative (SRQ and ERJ), the Clare Boothe Luce Foundation (ERJ), and the National Science Foundation through the Physics Frontier Center for Living Systems (PHY-2317138) (ERJ and NR).

