## Supplementary Information for "Spatially-structured inflammatory response in the presence of a uniform stimulus"

### Supporting Information Text

#### Materials and Methods.

**Dissociation protocol for scRNAseq screen.** We used the following buffers during dissociation: Euthanasia: .04% w/v Tricaine (Pentair) in E3 (1). Digestion: Final concentration .12 mg/mL Liberase-TL (Millipore Sigma, 5401020001) and 22.5 U/mL DNAse I (Worthington, LS006331) in .25% Trypsin/EDTA (Gibco, 25200-056). Mincing: Final concentration 58.25 U/mL DNAse I in PBS. Wash/quench: Final concentration .1% BSA in PBS. Larvae were held for 10 min in euthanasia buffer on ice, and then decapitated with a sterile razor blade. The body was transferred to scant mince buffer on a sterile petri dish and minced. Collected tissue was transferred to 300 uL of digestion buffer in an eppendorf tube. The tube was placed on a heat block at 30 C for 6 minutes, and a P1000 pipette was used to triturate. The samples were centrifuged at 400 g for 5 minutes and the digestion buffer was removed. Cells were resuspended in 1 mL wash buffer per tube; filtered through a 40 um cell strainer; centrifuged for 5 minutes at 400 g; and resuspended in 30-50 ul wash buffer. Cell density was estimated by counting on a hemocytometer prior to loading into a 10X Genomics lane.

**Null models for gene expression variability analysis.** We constructed null models for each sample and gene to capture the expected level of variability for the case where cells express the gene uniformly, but there is measured variability due to finite numbers of spots and uncertainties in cell boundaries, which creates variation in the true number of cells per cell region.

Let $g(n)$ be the probability of a cell region having $n$ cells, and $b(k|n)$ be the probability of observing $k$ counts in a region with $n$ cells. The null model is then $p(k) = \int \mathrm{d}n\, g(n) b(k|n)$. To model the distribution of counts per cell at a given total expression level, we take $b(k|n) = \mathrm{Binomial}(N, p)$, where $N$ is the total number of spots for the sample and gene, and $p = \frac{1}{n_c} \frac{n}{\bar{n}}$, where $n_c$ is the number of cell regions, and $n$ is the average number of cells per cell region, $\bar{n} = \int \mathrm{d}n\, n g(n)$. The effective true number of cells per cell region can vary either because a cell got fractionally subsected in segmentation (fewer than 1 cell/region), or because two cells from different layers overlap (more than 1 cell/region). Since we do not have access to the true distribution, we take a near-worst-case scenario where $g(n) \sim U(n_l, n_u)$: $g(n)$ is uniformly distributed between $n_l$ and $n_u$ cells/region. For Figure 2B, we take $n_l = .5$, $n_u = 2.5$; for Figure S3, we varied the width of the uniform distribution, $n_u - n_l$, between .5 and 4.5. We estimated the convolution integrals via a sum. To compare the variability in the data vs. the null model, we measured the ratio of the standard deviation of counts in the data and the null model, for each sample and gene.

**Spectral Graph Decomposition.** We define a graph of each tailfin sample, where each cell region is a node, and cells that are neighbors (share any boundary pixels) are connected by edges. The graph is unweighted. The adjacency matrix $A$ for the graph is:

$$A_{ij} = \begin{cases} 1 & \text{i,j neighbors} \\ 0 & \text{Otherwise.} \end{cases} \quad [1]$$

Note that $A$ is symmetric $A^T = A$. Letting the degree matrix $D$ be:

$$D_{ij} = \begin{cases} 1 + \sum_j A_{ij} & \text{i=j} \\ 0 & \text{Otherwise.} \end{cases} \quad [2]$$

The degree matrix diagonal elements $D_{ii}$ count how many edges the node $i$ is a part of. We define the normalized adjacency matrix $\hat{A}$ as:

$$\hat{A} = (D^+)^{1/2} A (D^+)^{1/2}, \quad [3]$$

where the inverse degree matrix $D^+$ is:

$$D^+_{ij} = \begin{cases} \frac{1}{D_{ij}} & \text{i=j} \\ 0 & \text{Otherwise.} \end{cases} \quad [4]$$

Finally, the symmetric normalized graph Laplacian matrix is:

$$L = I - \hat{A}. \quad [5]$$

The graph laplacian $L$ is so-named because it is a 'discrete' version of the standard Laplacian differential operator on the graph. Since $L$ is symmetric, it has real eigenvalues and an orthonormal basis of eigenvectors which can be computed through the eigendecomposition of $\hat{A}$ since $\hat{A}$ and $L$ commute:

$$\hat{A} = V \Lambda V^T, \quad [6]$$

where $\Lambda = diag[\lambda_1, ...\lambda_n]$ is the matrix of eigenvalues of $\hat{A}$, ordered from high to low. The eigenvectors are then the columns of $V$: $V = [\vec{v}_1, ...\vec{v}_n]$.

Note that $n$ is the number of cells in the sample, and the eigenvectors also have length $n$.

For any quantity measured on each cell, $\vec{s}$, we can compute the projection onto spatial mode $j$ as:

$$\vec{p}_j = (\vec{s} \cdot \vec{v}_j)\vec{v}_j, \quad [7]$$

where the coefficients $a_j = \vec{s} \cdot \vec{v}_j$ are analogous to Fourier coefficients. Thus the power corresponding to mode $j$ is $a_j^2$.

**Coons patch bilinear interpolation.** The Coons patch (2) is defined as follows: given four boundary curves, $c_0(s)$, $c_1(s)$, $d_0(t)$, $d_1(t)$, which meet at the four corners $c_0(0) = d_0(0)$, $c_0(1) = d_1(0)$, $c_1(0) = d_0(1)$, $c_1(1) = d_1(1)$, the linear interpolation between $c_0$ and $c_1$ is:

$$L_c(s,t) = (1-t)c_0(s) + tc_1(s), \qquad [8]$$

and the linear interpolation between $d_0$ and $d_1$ is:

$$L_d(s,t) = (1-s)d_0(t) + sd_1(t). \qquad [9]$$

We further define:

$$B(s,t) = c_0(0)(1-s)(1-t) + c_0(1)s(1-t) + c_1(0)(1-s)t + c_1(1)st. \qquad [10]$$

Then the Coons patch is the surface parametrized by $s,t$ such that

$$C(s,t) = L_c(s,t) + L_d(s,t) - B(s,t). \qquad [11]$$

We defined boundary curves for each sample by detecting corners on mask images via the corner harris function in skimage.feature, and used a 1d cubic spline interpolation to transform the list of pixels between corners into boundary curves. We used these curves to define the Coons patch, as specified above. To map cell centroid coordinates $(x,y)$ from the original space to their corresponding coordinates $(s,t)$ in the Coons patch, we computed a lookup table for the mapping over a fine grid (400x100, axial x outer-inner), and mapped to the nearest grid point.

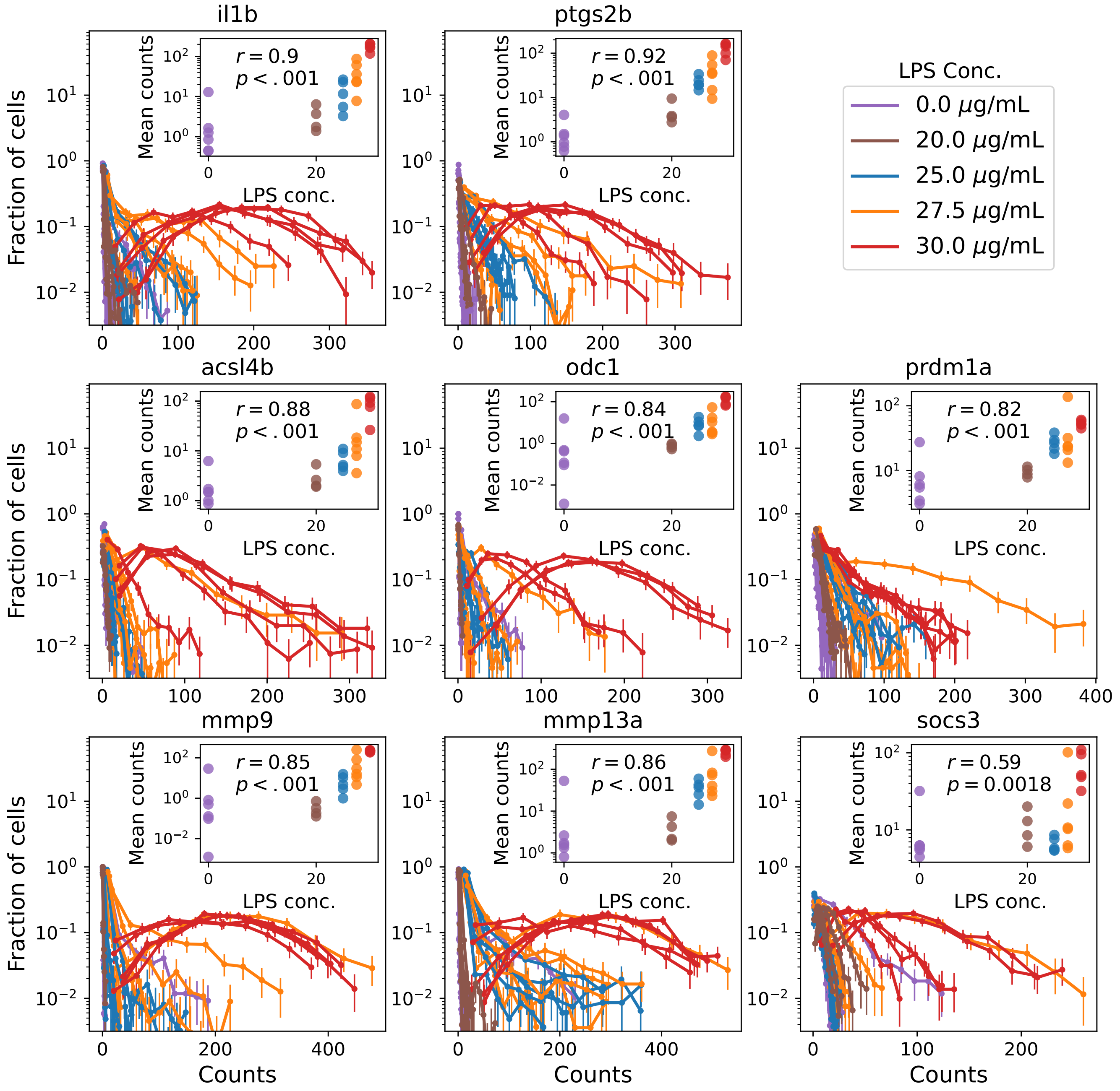


**Fig. S1.** Same as Figure 1B., with spot count as the measure of gene expression. Distributions of counts across cell regions in each sample. Insets: average counts and Spearman correlation coefficient; significance of the correlation via a permutation test.

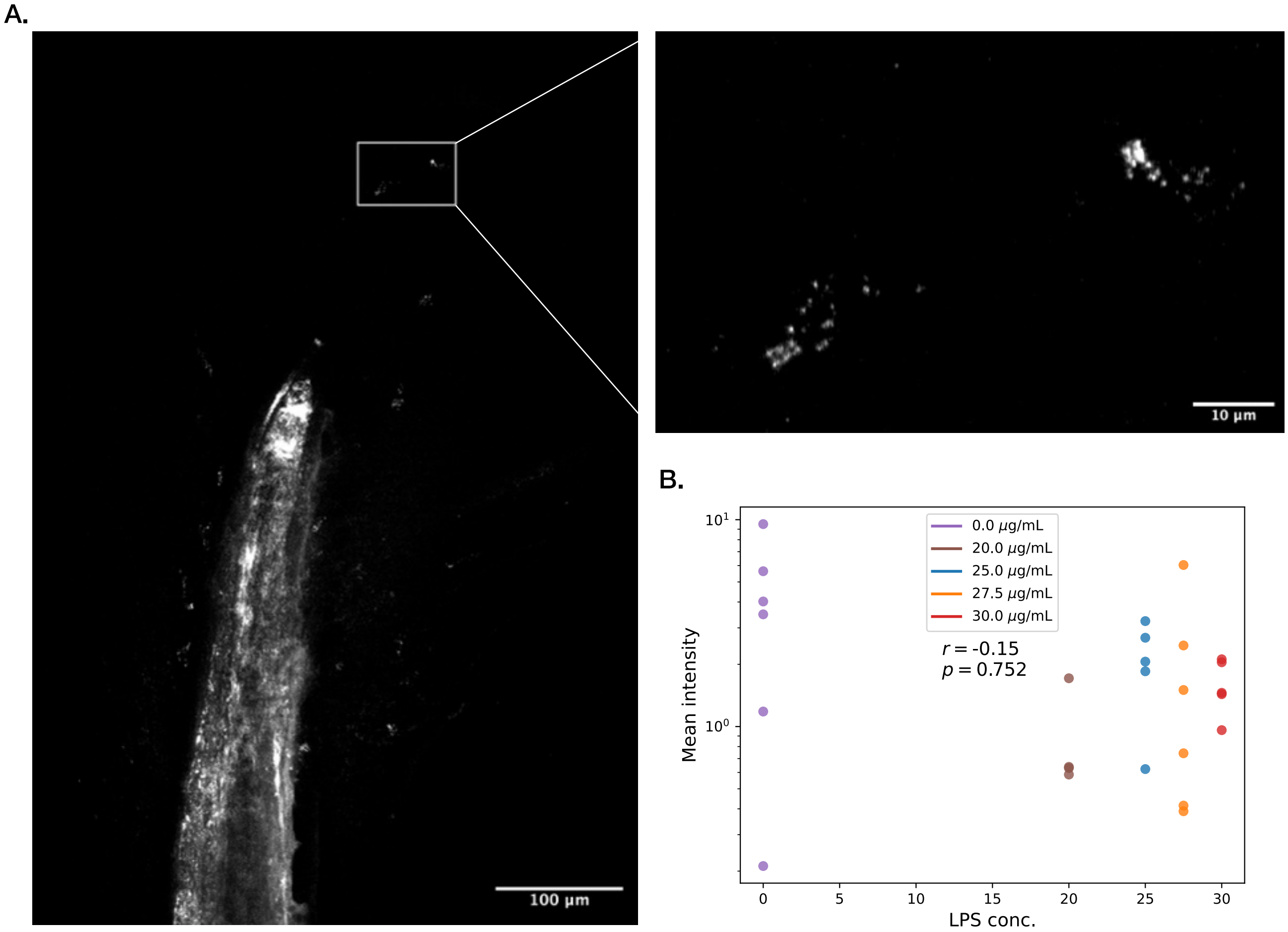


**Fig. S2.** A. Example of labeling of the macrophage marker mpeg1.1 in a tailfin sample. The isolated small labeled regions are consistent with isolated macrophages. ROI–two putative macrophages. B. Average intensity per segmented cell region for the mpeg1.1 gene channel across all samples shows no trend.

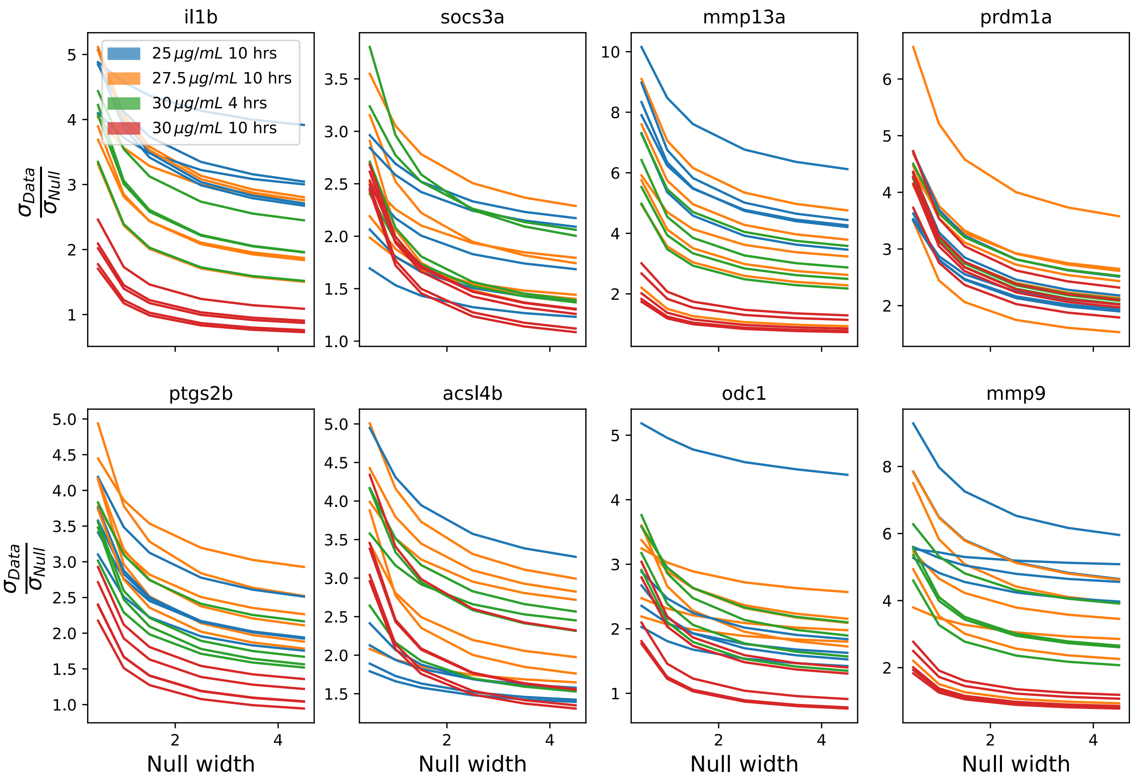


**Fig. S3.** Ratio of widths of count distributions from data vs. null, under a range of assumptions of the true number of cells per segmented cell region. In each case, the null distribution assumes a uniform distribution of cells per cell region. The ratio declines with increased uncertainty about the true number of cells per cell region, but remains high even for broad distributions.

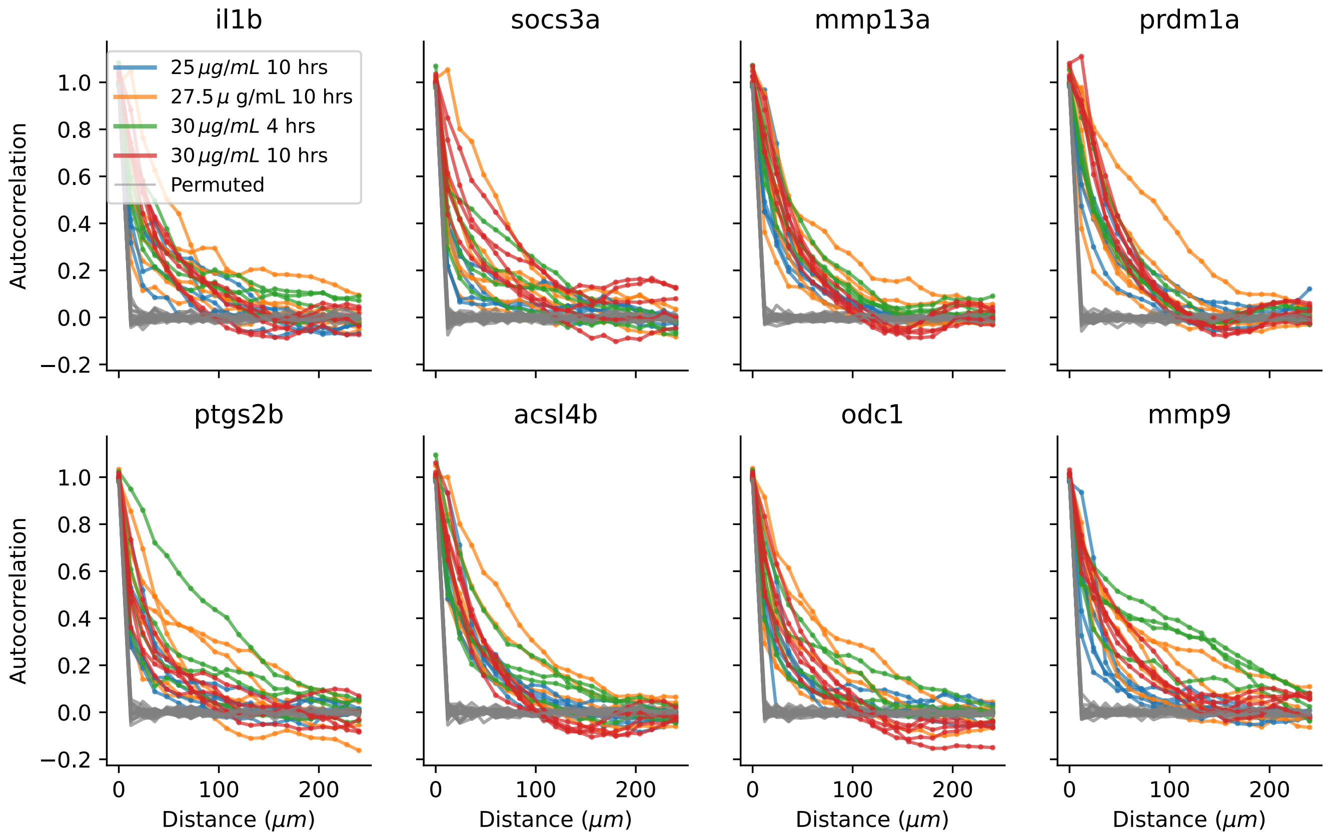


**Fig. S4.** Spatial autocorrelations across all genes, as in Figure 2B.

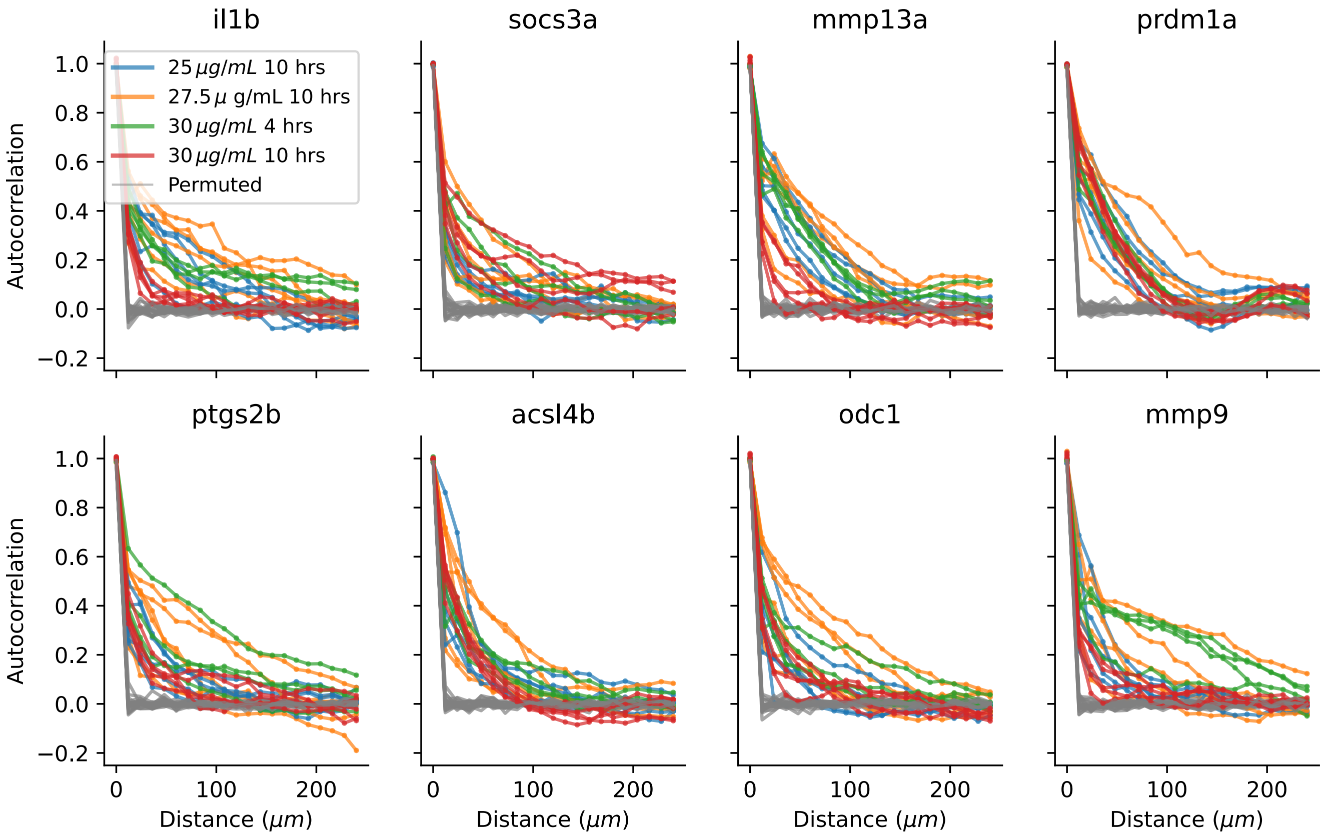


**Fig. S5.** Spatial autocorrelations across all genes, with gene expression measured via spot counts.

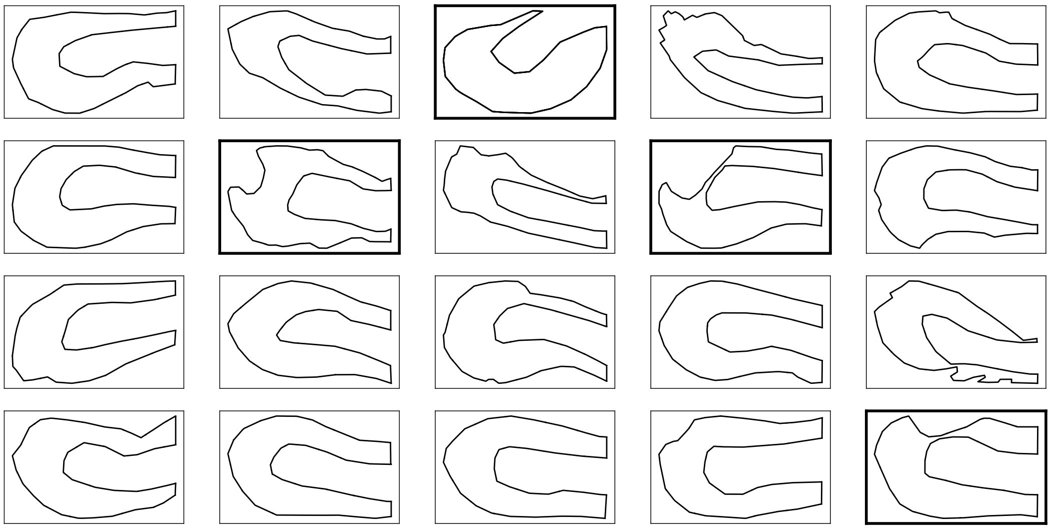

**Fig. S6.** Outlines of all tail samples, with bold borders indicating samples not used in the rectangle-mapping analysis. We note that in the interests of including as many samples as possible, we included those with rough boundaries.

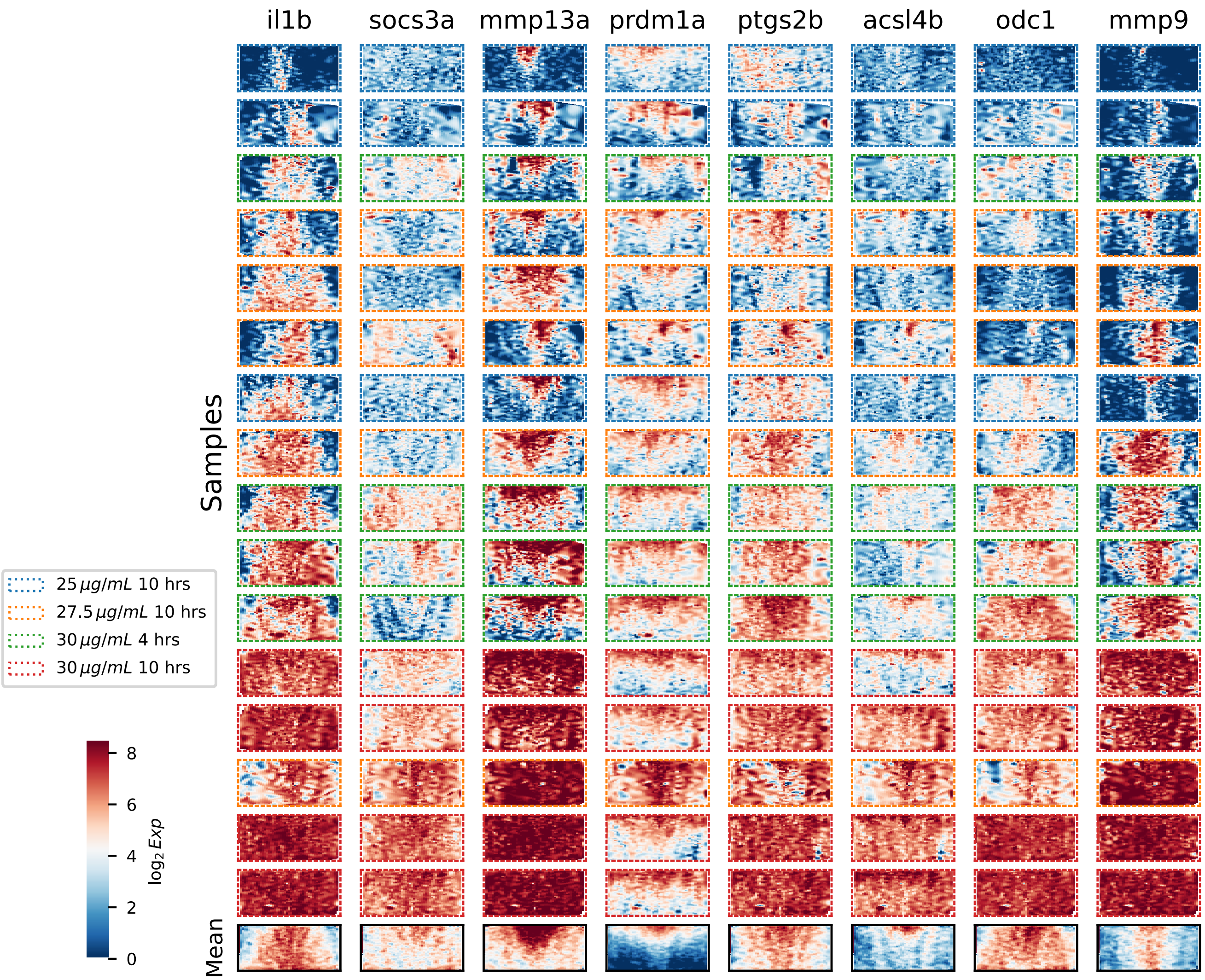


**Fig. S7.** Same as Figure 3B, with spot counts as the measure of gene expression.

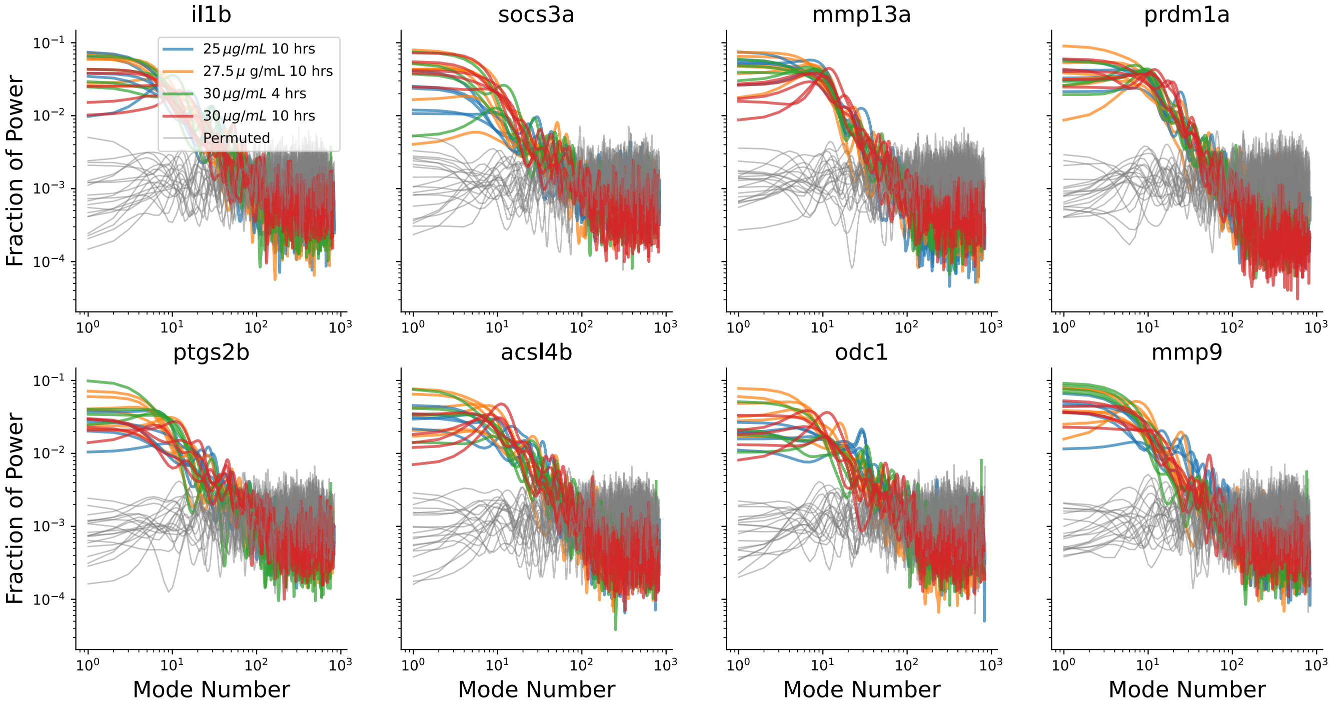


**Fig. S8.** Fraction of power per graph mode, as in Figure 4B, for all genes and samples.

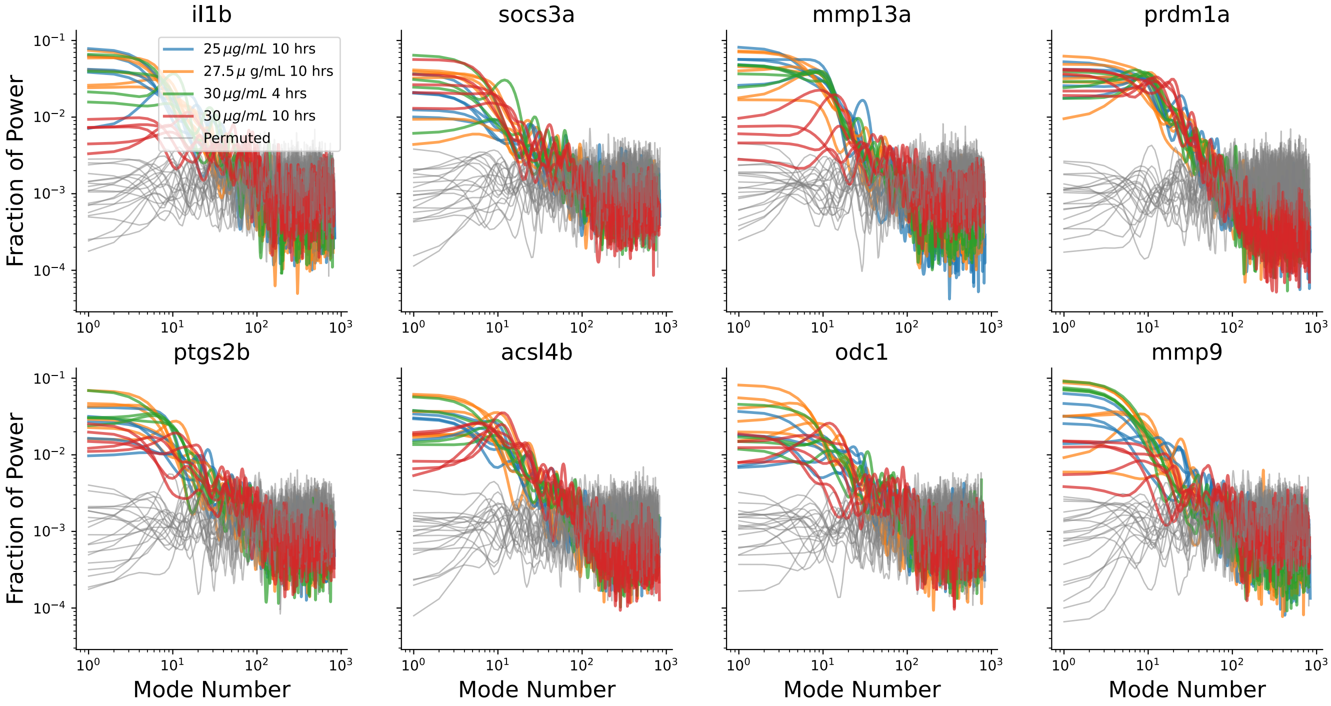


**Fig. S9.** Fraction of power per graph mode, as in Figure 4B, with spot counts as the measure of gene expression.

 

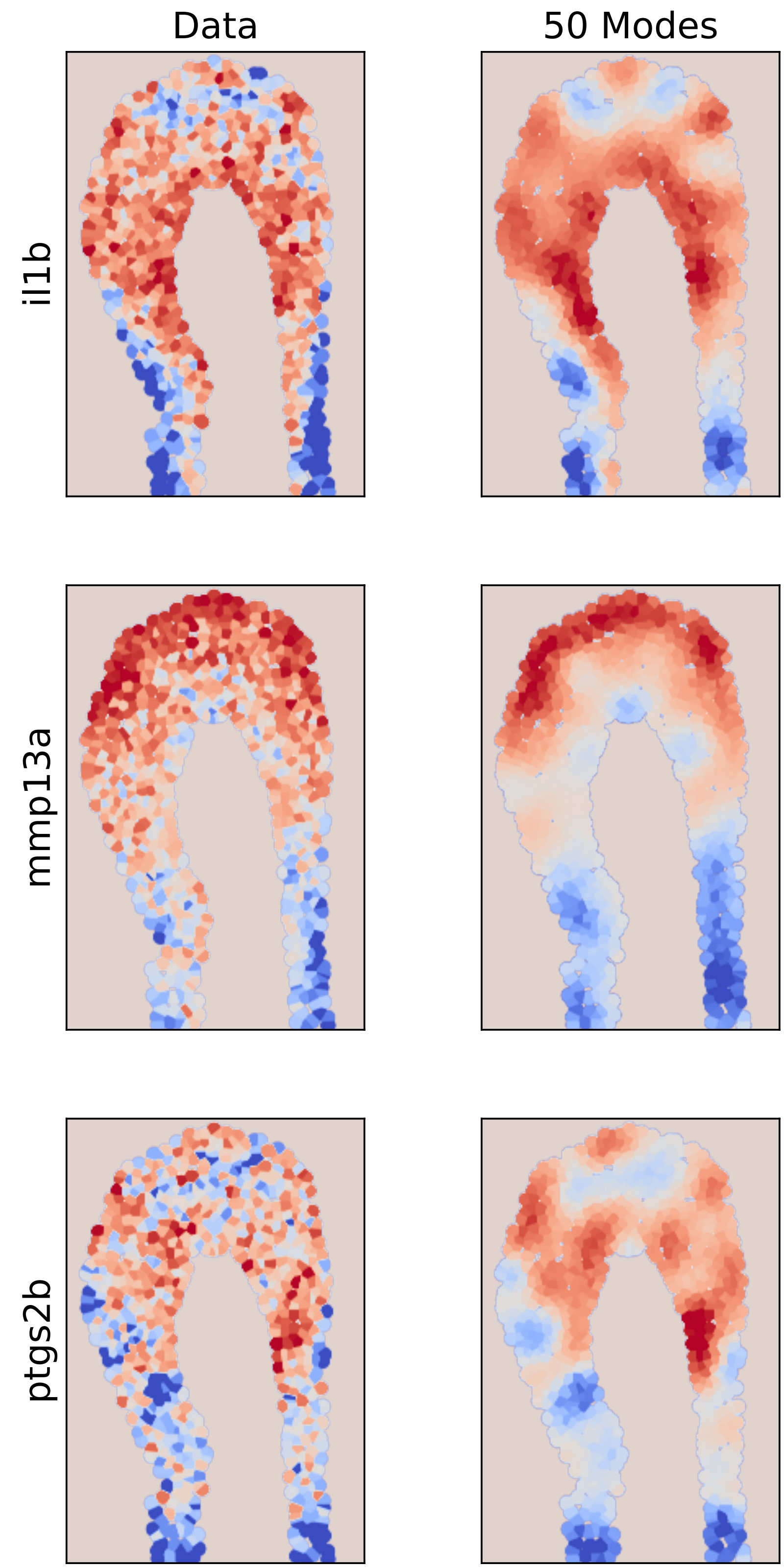


**Fig. S10.** Same as Figure 4C, with spot counts as the measure of gene expression.

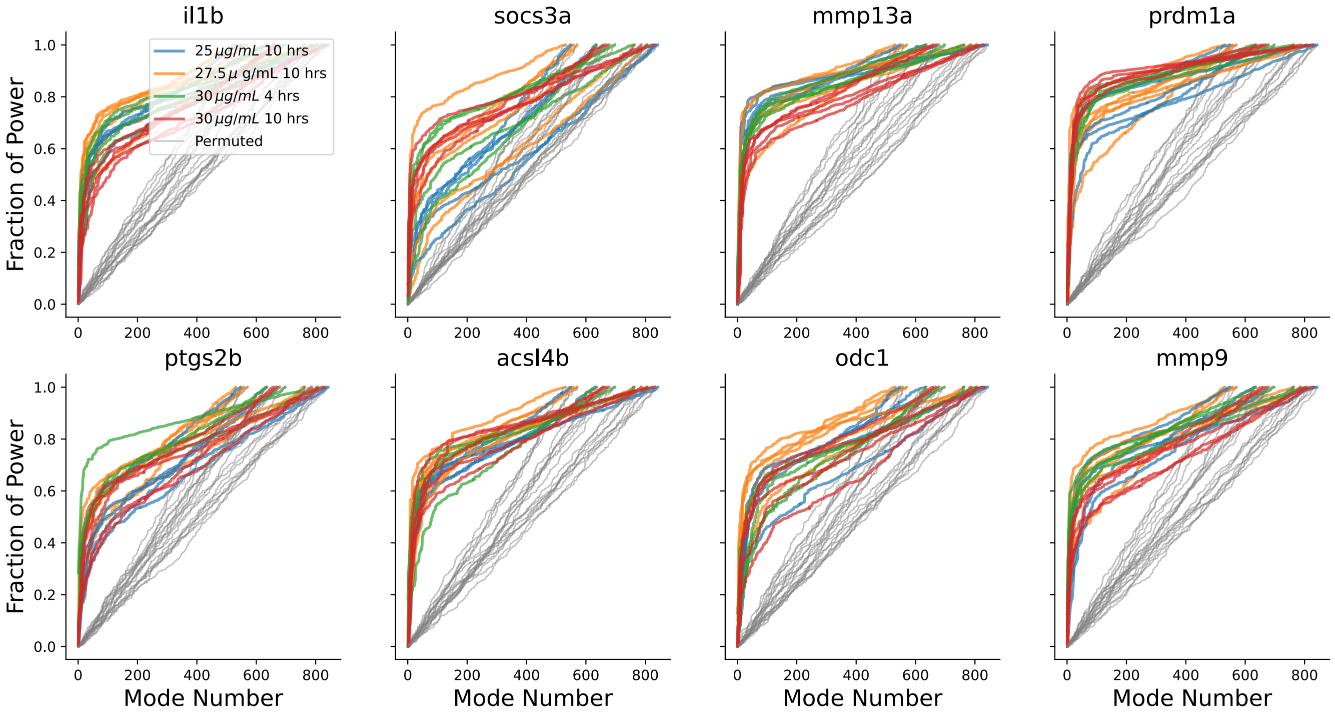


**Fig. S11.** Fraction of the variance explained by modes up to the given number, for all genes.

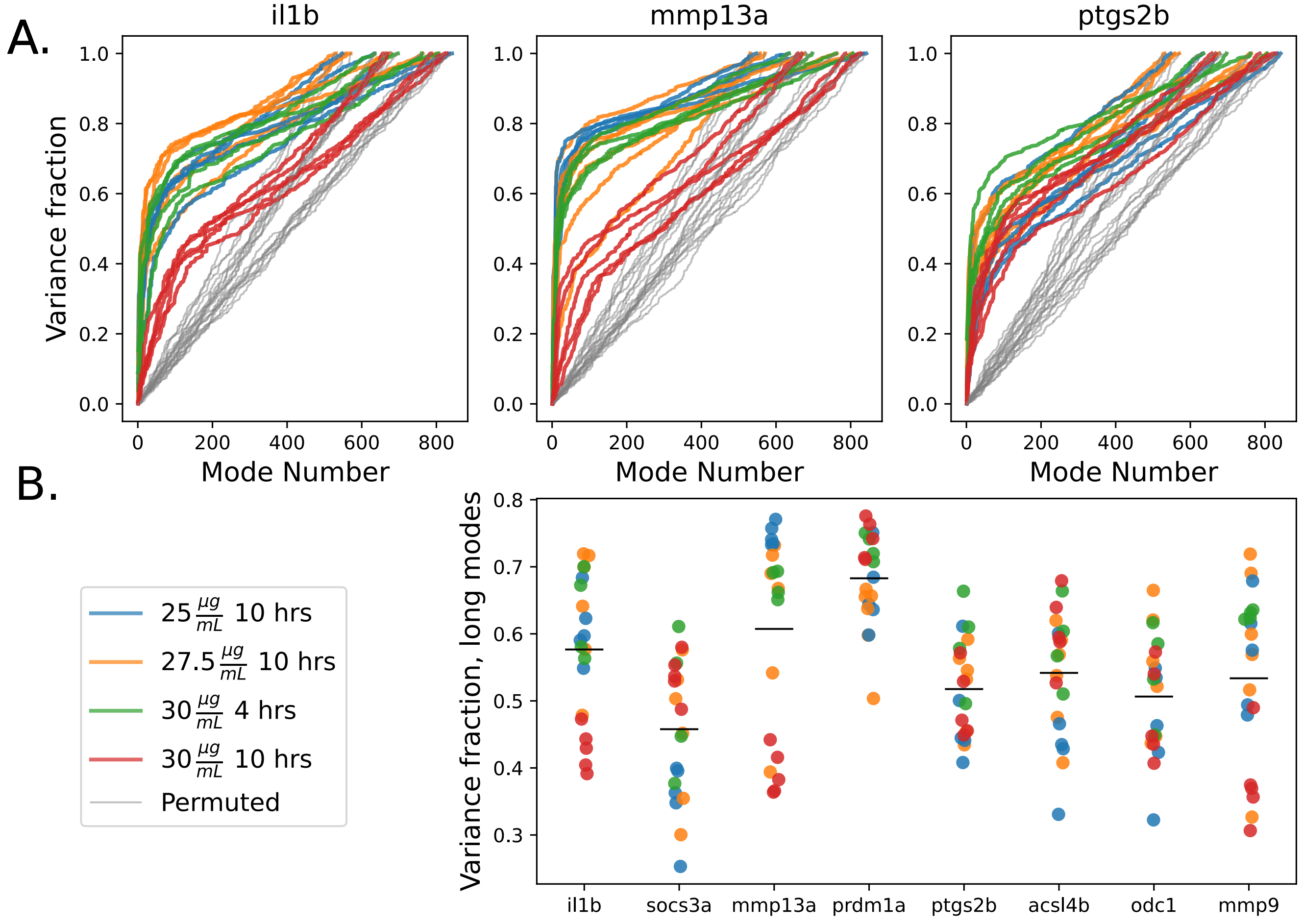


**Fig. S12.** Same as Figure 5B, with spot counts as the measure of gene expression.
